# Sex-specific decline in prefrontal cortex mitochondrial bioenergetics in aging baboons correlates with walking speed

**DOI:** 10.1101/2024.09.19.613684

**Authors:** Daniel A. Adekunbi, Hillary F. Huber, Gloria A. Benavides, Ran Tian, Cun Li, Peter W. Nathanielsz, Jianhua Zhang, Victor Darley-Usmar, Laura A. Cox, Adam B. Salmon

## Abstract

Mitochondria play a crucial role in brain aging due to their involvement in bioenergetics, neuroinflammation and brain steroid synthesis. Mitochondrial dysfunction is linked to age-related neurodegenerative diseases, including Alzheimer’s disease and Parkinson’s disease. We investigated changes in the activities of the electron transport chain (ETC) complexes in normally aging baboon brains and determined how these changes relate to donor sex, morning cortisol levels, and walking speed. Using a novel approach, we assessed mitochondrial bioenergetics from frozen prefrontal cortex (PFC) tissues from a large cohort (60 individuals) of well-characterized aging baboons (6.6–22.8 years, approximately equivalent to 26.4–91.2 human years). Aging was associated with a decline in mitochondrial ETC complexes in the PFC, which was more pronounced when activities were normalized for citrate synthase activity, suggesting that the decline in respiration is predominantly driven by changes in the specific activity of individual complexes rather than changes in mitochondrial number. Moreover, when donor sex was used as a covariate, we found that mitochondrial respiration was preserved with age in females, whereas males showed significant loss of ETC activity with age. Males had higher activities of each individual ETC complex and greater lactate dehydrogenase activity relative to females. Circulating cortisol levels correlated only with complex II-linked respiration in males. We also observed a robust positive predictive relationship between walking speed and respiration linked to complexes I, III, and IV in males but not in females. This data reveals a previously unknown link between aging and bioenergetics across multiple tissues linking frailty and bioenergetic function. This study highlights a potential molecular mechanism for sexual dimorphism in brain resilience and suggests that in males changes in PFC bioenergetics contribute to reduced motor function with age.

## 1.0 Introduction

Aging leads to a progressive decline in cognitive function and is a major risk factor for neurodegenerative diseases (Niccoli and Patridge, 2012). Age-related cognitive decline is linked to decreased brain volume, potentially arising from reduced neuronal size and synaptic density (Terry and Katzman, 2001; Harada et al., 2013; Resnick et al., 2003). Reduced oligodendrocyte numbers lead to a loss of myelination, which impairs neural signal transmission (Sams, 2021). Activation of microglia during aging also increases brain inflammation, further exacerbating cognitive decline (Dheen et al., 2007). Mitochondria play a central role in these processes due to their contribution to energy production and brain steroid synthesis (Lejri et al., 2018). Mitochondrial dysfunction has been linked to age-related neurodegenerative diseases, including Alzheimer’s and Parkinson’s diseases (Wang et al., 2020; Weidling et al., 2020; Austad et al., 2021; Misrani et al., 2021). Age-associated changes in ATP production, reactive oxygen species (ROS) production, and mitochondrial dynamics have all been reported as potential drivers of altered neuronal signaling, synaptic plasticity, and impaired memory formation (Yin et al., 2012). However, there are significant challenges in identifying the proximal causes of these impairments in normally aging brains, and the extent to which these involve mitochondrial electron transport chain (ETC) dysregulation is incompletely understood.

The multi-enzyme complexes that make up the components of oxidative phosphorylation include complexes I-IV, cytochrome c, and the ATP synthase and the small molecule electron carrier ubiquinone which together control ATP production (Papa et al., 2012). About 90% of inspired oxygen is consumed by mitochondria through complex IV of the ETC as part of oxidative phosphorylation to produce ATP. In the brain, high energy consumption is required for the synthesis of neurotransmitters and maintenance of membrane potential for action potential generation and synaptic signaling. The effects of age on brain ETC complex activities are limited and contradictory. For example, the activity of cytochrome c oxidase (complex IV) increases with age in the mouse cerebral cortex, while complexes I, II, and III are unaffected (Sharman and Bondy, 2001). In another mouse study using whole brain mitochondrial preparations, activities of complexes II and III decreased in older animals, while complexes I and IV were unchanged (Kwong and Sohal, 2000). In a rat study, complex I activity in the cerebral cortex decreased with age, while other complex activities were unaffected (Cocco et al., 2005). While there seems to be no uniformity in changes to respiratory chain activities during aging, it appears that brain region and species specificity influence the observed changes in mitochondrial complex activities during aging.

Further, there are challenges in reconciling molecular and functional changes with age in rodent brains with those that occur in humans and other primates. There are practical limitations in accessing human brain samples that are not confounded by diseases or trauma across the life course. Even with opportunistic collection of human postmortem brain tissues, the interval of tissue collection before preservation can take several hours, during which protein quality may be compromised. The need to assess mitochondrial respiration, the gold standard measurement of mitochondrial function, in freshly collected samples further limits the availability of brain data in human subjects. One human study using brain samples from traffic accident patients reported a tendency for mitochondrial complex I activity to decrease with age in the hippocampus and frontal cortex (Venkateshappa et al., 2012). However, the study had a relatively small sample size (n=17-24), did not account for sex effects despite the role of sex hormones in brain mitochondrial function (Yao et al., 2011), and did not examine other mitochondrial features and complex activities. The use of appropriate experimental models that share similar brain homology with humans can address some of these gaps.

Cognitive decline with age shows similarities between humans and multiple nonhuman primate (NHP) species (Lacreuse et al., 2020). Baboons, in particular, share closer physiology and genetics with humans compared to other commonly used experimental NHP models (Cox et al., 2013). Evidence of the natural occurrence of age-related cognitive decline in baboons further supports their use for translational studies to humans (Lizarraga et al., 2020). The prefrontal cortex (PFC) is one of the key brain regions most affected by age (Raz et al., 2005) and is central to cognitive abilities such as decision-making, problem-solving, working memory, temporal processing, and goal-oriented behavior (Friedman et al., 2022). It is also part of the executive locomotor pathway (Hamacher et al., 2015; Nóbrega-Sousa et al., 2020), with specific cognitive capabilities such as visuospatial skills, cognitive processing speed, and attention required for motor tasks like walking. Walking speed is one of the best single measurement clinical predictors of morbidity and mortality (Abellan van Kan et al., 2009, Studenski et al., 2011). In older adults, incremental decline in gait speed is associated with poorer executive function (Atkinson et al., 2007), indicating that deficits in executive function may contribute to gait impairment. However, the processes underlying this interaction remain unclear. We previously demonstrated that aging results in a decline in walking speed, a marker of frailty, in male and female baboons (Huber et al., 2021). Whether age-related changes in energy metabolism in the PFC can be predicted by walking speed has not been considered.

Additionally, several studies have suggested that increased cortisol levels are associated with decreased cognitive abilities (Lupien et al., 2004; Li et al., 2006; Lee et al., 2007). This relationship, however, is complicated by the timing of cortisol measurement, participants’ disease state, and type of cognitive assessment. Glucocorticoids (cortisol in humans and corticosterone in rodents) and other steroids are produced and metabolized by mitochondria (Bose et al., 2002). Reciprocally, mitochondria are sensitive to glucocorticoids (GC), and the presence of GC receptors in mitochondria suggests direct actions in this organelle (Psarra and Sekeris, 2009). GC exert biphasic effects in neural tissues (Choi and Han, 2021; Du et al., 2009). Acute stress increases mitochondrial activity within the brain in a process that utilizes glucose and oxygen to provide energy for an appropriate physiological response and stress adaptation (Picard et al., 2018). By contrast, chronic exposure to high GC levels induces abnormal mitochondrial morphology, dysregulates mitochondrial fusion and fission processes, and inhibits mitochondrial trafficking to synaptic regions in response to energy demand, culminating in brain atrophy (Choi and Han, 2021; Kiryu-Seo et al., 2016). It is not clear if there is a relationship between the effect of age on circulating cortisol and mitochondrial activity in the brain.

Building on our previous studies, which demonstrated an age-dependent decline in cortisol concentrations in female baboons (Yang et al., 2017; Nathanielsz et al., 2020) we have integrated these findings into a bioenergetic profile. We examined the effect of age on circulating cortisol concentrations in a larger sample size that includes both male and female baboons and determined the corresponding changes in PFC mitochondrial ETC activities, mitochondrial content, and lactate dehydrogenase activity across the life course. Additionally, we evaluated the relationship between circulating cortisol and PFC mitochondrial ETC activity. Importantly, we revealed a novel and intriguing result which shows a robust association of PFC mitochondrial ETC activity with walking speed at the individual animal level across the lifespan.

## 2.0 Methods

### 2.1. Animals

The Institutional Animal Care and Use Committee of Texas Biomedical Research Institute (TX Biomed) approved all procedures involving animals. The animal facilities at the Southwest National Primate Research Center (SNPRC) within the campus of TX Biomed are fully accredited by the Association for Assessment and Accreditation of Laboratory Animal Care International (AAALAC) and adhere to the guidelines of the National Institutes of Health (NIH) and the U.S. Department of Agriculture.

Baboons (*Papio* species) were housed and maintained in outdoor social groups and fed *ad libitum* with normal monkey diet (Purina Monkey Diet 5LEO; 2.98 kcal/g). The welfare of the animals was enhanced by providing enrichment, such as toys, food treats, and music, which were offered daily under the supervision of the veterinary and behavioral staff at SNPRC. All baboons received veterinarian physical examinations at least once per year and were healthy at the time of study. Animals were aged between 6.6 and 22.8 years (approximate human equivalent, 26.4 – 91.2 years), n=60. The baboons were part of a cohort of animals studied throughout the life course to identify and establish effects of physiological aging (Huber et al., 2021; Kuo et al., 2018; Huber et al., 2015; Cox et al., 2023; Grilo et al., 2024).

### 2.2. Sample collection

Healthy male (6.6 – 22.8 years, n= 26) and female (7.4 – 22.1 years, n=34) baboons were tranquilized with ketamine hydrochloride (7-10 mg/kg intramuscular injection) after an overnight fast. Blood was collected to obtain serum for determination of circulating cortisol concentrations. Parts of the female cortisol data were published in a separate study (Yang et al., 2017). Here, we increased female cortisol data and added male data. We previously demonstrated that the tranquilizing dose of ketamine used for immobilization does not raise circulating cortisol in the 10 min maximum time needed to take a blood sample (Yang et al., 2017). Blood for cortisol measurement was obtained 2-3 days before necropsy. Baboons were euthanized by exsanguination while under general anesthesia (1-2% isoflurane), as approved by the American Veterinary Medical Association. We have shown that euthanasia by intravenous agents can alter tissue structure (Grieves et al., 2008). Following cardiac asystole, brains were collected according to a standardized protocol about 30 min from time of death, between 8:00-10:00 AM to minimize potential variation from circadian rhythms. Brain tissues were collected under aseptic sterile conditions and the PFC was dissected from the left side of the brain. About 3 mm deep samples of gray matter from the PFC were collected, extending from the posterior part of the precentral sulcus to the intersection of the precentral sulcus and lateral sulcus.

### 2.3. Hormone quantification

Serum cortisol concentration was quantified as we previously reported (Yang et al., 2017) using a chemiluminescent immunoassay on an Immulite 1000 analyzer with kits from Siemens Healthcare Diagnostics (Flanders, NJ, USA). The intraassay and interassay coefficients of variation for cortisol were 5.6 and 8.4, respectively.

### 2.4. Bioenergetic Measurements

Mitochondrial electron transport chain (ETC) activity of baboon PFC across the life course was determined using a previously described respirometry assay for frozen tissues (Acin-Perez et al., 2020; Benavides et al., 2022). The approach involves supplementing cytochrome c lost in cryopreserved mitochondria to complete the flow of the ETC, enabling a well-controlled measurement of maximal oxygen consumption of the ETC using physiological electron donors and acceptors and has been applied to the measurement of mitochondrial ETC in human brain (Benavides et al., 2022). Following baboon tissue optimization, we identified 6 µg protein concentration as optimal for the respirometry assay. Pulverized PFC samples were homogenized in mitochondrial assay buffer; MAS containing 70 mM sucrose, 220 mM mannitol, 5 mM KH_2_PO_4_, 5 mM MgCl_2_, 1 mM EGTA, 2 mM HEPES, using mortar and pestle over liquid nitrogen. Protein concentration was determined by DC Lowry protein assay. PFC homogenates were loaded onto a 96-well cell plate, centrifuge at 2000x*g* for 20 min at 4°C, followed by serial injection of mitochondrial substrates and modulators using the Agilent XF96 Extracellular Flux Analyzer (North Billerica, MA, USA). Respirometry measurements were performed to evaluate the activity of mitochondrial ETC complexes I, II, III and IV. The following substrates were added to the seahorse cartridges to stimulate complex activities: Complex I-reduced nicotinamide adenine dinucleotide (NADH; 1 mM), Complex II-combination of succinate (10 mM) and rotenone (2 µM), Complex III – Duroquinol (2.5 mM), and Complex IV – tetra methyl phenylene diamine, TMPD (0.5 mM) and ascorbic acid (2 mM) cocktail. Mitochondrial complexes were inhibited with 2 µM rotenone, 10 µM Antimycin A, or 20 mM azide. Oxygen consumption rate (OCR) data following the seahorse runs were extracted using the Agilent Wave analysis software. The activities of the complexes were calculated as follow: Complex I-post-NADH injection OCR minus post rotenone injection OCR. Complex II-post-succinate and rotenone cocktail injection OCR minus post-antimycin A injection OCR. Complex III-post-duroquinol injection OCR minus post-antimycin A injection OCR. Complex IV-post-TMPD and ascorbic acid cocktail OCR injection minus post-azide injection OCR. OCR values of each mitochondrial complex were normalized to protein.

### 2.5. Citrate synthase and lactate dehydrogenase activity assays

The reaction mixture for citrate synthase (CS) assay contains 1 mM dithiobis-(2-nitobenzoic acid) (DNTB), 0.2 mM acetyl CoA, 0.1M Tris-HCl and 10mM Oxaloacetate at pH 8.1. PFC homogenates were added to the reaction mixture in triplicates. Enzyme activity was determined by spectrophotometry and absorbance captured at 412 nm for 5 min. CS activity was calculated as nmol/min/mg protein from the change of absorbance. For measurement of lactate dehydrogenase (LDH) activity, NADH and pyruvate were incubated with protein lysates. We then measured the disappearance of NADH absorption at 340 nM and calculated the activity as nmol/min/mg protein.

### 2.6. Walking speed

Walking speed data were part of previously published studies (Huber et al., 2021; Huber et al., 2015) where we demonstrated age-dependent decline in walking speed in both male and female baboons. These data are used for correlative analysis with corresponding PFC bioenergetic data in an individual animal to determine potential interaction between brain energy metabolism and locomotor changes with age. About 2 years prior to necropsy, behavioral observations of walking speed were conducted under ad libitum conditions between 8:00–11:00am by timing with a stopwatch how long it took baboons to walk between landmarks within home cages (Huber et al., 2021; Huber et al., 2015). We only recorded straight walks from one landmark to another for a distance of at least 2 meters. Instances of walking toward food and social interactions were not recorded to avoid bias by underlying motivation. For each subject, 5–15 bouts of walking were timed and recorded over a single 3-hour period, then averaged. We have shown that outdoor temperature and humidity do not affect walking speed (Huber et al., 2021). Walking speed was calculated as distance (cm)/time (sec).

### 2.7. Statistical analysis

To determine the effects of age on serum cortisol concentrations, the activities of each mitochondrial complex, CS, and LDH were regressed against age across the life course (males, 6.6 – 22.8 years, and females, 7.4 – 22.1 years) using a simple linear regression. Walking speed and serum cortisol concentrations in an individual animal were correlated with the PFC mitochondrial features using Pearson’s correlation method. Comparison of mitochondrial complex activities between male and female, independent of donor age was achieved by Student’s t-test. Significance was set at p < 0.05.

## 3.0 Results

### 3.1. Aging decreases mitochondrial ETC activity in the baboon prefrontal cortex

The bivariate correlation of age with each ETC complex activity (measured as OCR) showed age-related decline in respiration with complex I (p=0.014), II (p=0.011), and III (p=0.017) in the baboon PFC when male and female data were combined (Fig 1 A-C). Respiration through complex IV tended to decline with age (p=0.086; Fig 1D). CS is an enzyme located in the mitochondrial matrix which is not a component of oxidative phosphorylation and is typically used as a surrogate for the amount of mitochondria in a tissue. There was no significant association between aging and the CS activity (Figure 1E). LDH is a cytosolic enzyme involved in glycolysis and similarly did not show any significant association with age (Figure 1F). Importantly, the CS/LDH ratio does not show any significant correlation with age which suggest there is not an age dependent bias in the preparation of the homogenates. When we normalized each complex to CS activity, we observed a more robust age-related decline in respiratory activities of all the ETC complexes in combined data from male and female baboons (C-I/CS p=0.0014; C-II/CS p=0.0017; CIII/CS p=0.0024; CIV-IV p=0.0028; Fig 2 A-D). This suggests that there is a deficit in the specific activity of the mitochondrial ETC components with a minimal contribution from changes in mitochondrial number with age.

**Fig. 1:**
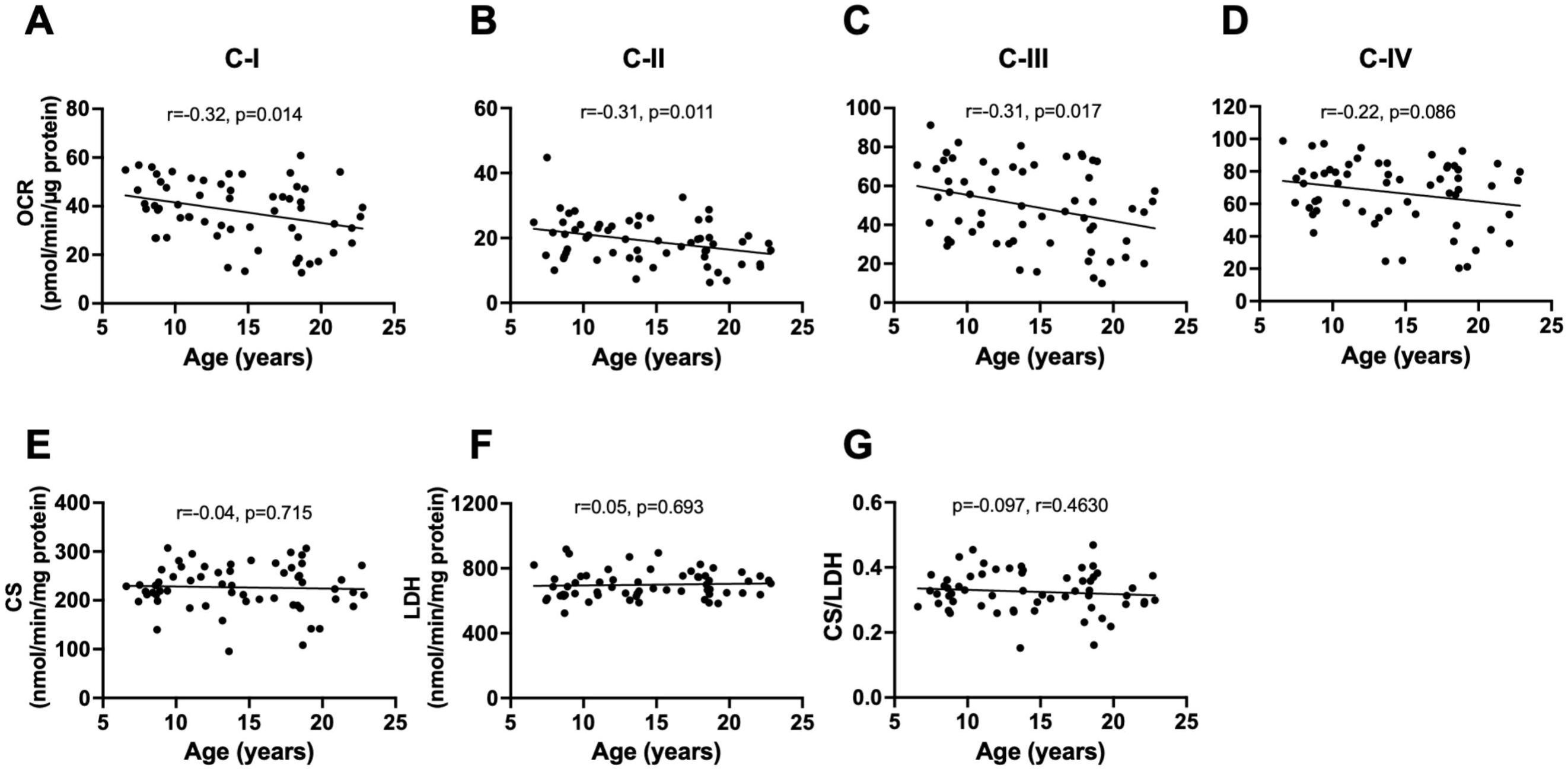
Linear regression of mitochondrial bioenergetics in the prefrontal cortex with age in male and female baboons. Data from male and female baboons were pooled. Plots of mitochondrial complexes activities: C-I (A), C-II (B), C-III (C), and C-IV (D) versus donor age. Citrate synthase (CS) activity (E), lactate dehydrogenase (LDH) activity (F), and CS to LDH ratio (G) were also regressed against age. The slope of the regression is represented by a thick line, and each individual dot represents data from an individual animal (n=60).

**Fig. 2:**
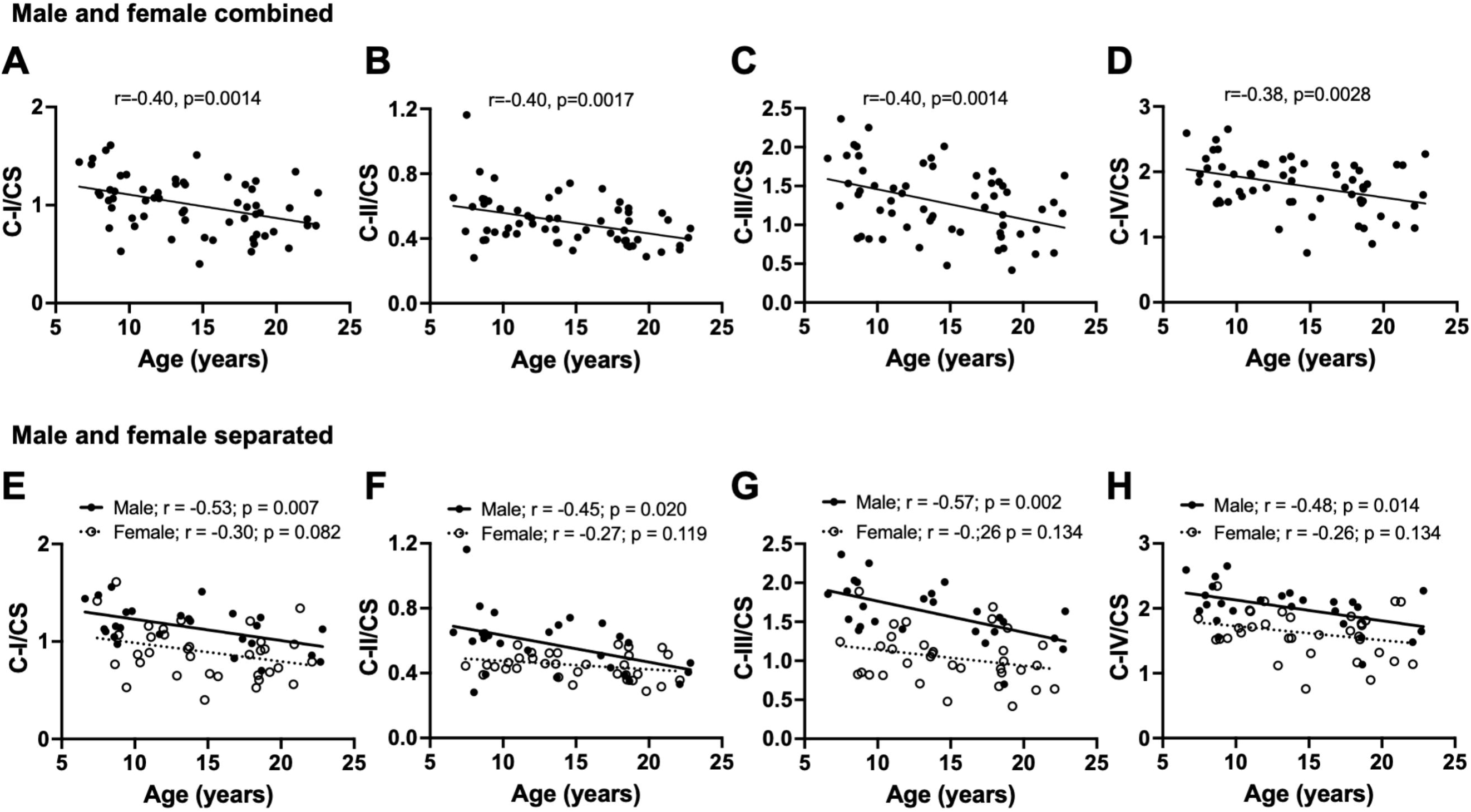
Linear regression of normalized mitochondrial complex activities in the prefrontal cortex with age in male and female baboons. Regression analysis was conducted on either combined male and female data or separated by sex. Plots of citrate synthase (CS)-normalized mitochondrial respiration due to complex I: C-I/CS (A), complex II: C-II/CS (B), complex III: C-III/CS (C), and complex IV: C-IV/CS (D) versus donor age when male and female data were combined (n=60). Linear regression of C-I/CS (E), C-II/CS (F), C-III/CS (G), and C-IV/CS (H) with age, showing separate analysis for males and females.

When we separately analyzed normalized complex activity by sex, only males showed a significant loss of complex-dependent respiration with age (C-I/CS p=0.007; C-II/CS p=0.020; CIII/CS p=0.002; CIV/CS p=0.014) while the decline in females was not significant (C-I/CS p=0.082; C-II/CS p=0.119; CIII/CS p=0.134; CIV/CS p=0.134; Fig 2 E-H). We also observed the expected positive relationship (p<0.001) between complexes I to IV-dependent respiration and CS activity in both male and female animals consistent with CS acting as a surrogate marker for mitochondrial amount (Supplemental Fig 1). It is important to note that since changes in CS activity is not associated with age (Figure 1E) the progressive decrease with ETC activities with age (Fig 1,2 are not strongly influenced by age-dependent changes in mitochondrial number.

### 3.2. Male baboons exhibit higher mitochondrial ETC activity than females

We next compared activities of mitochondrial ETC complexes (I to IV), CS, and LDH as well as CS/LDH ratio between male and female baboons, independent of age. Males had higher OCR via complex I, II, III, and IV in the PFC relative to females (C-I p=0.002; C-II p=0.008; C-III p<0.0001; CIV p=0.021; Fig 3 A-D). LDH activity was also higher in males compared to females (p=0.023; Fig 3 F), whereas CS activity and the CS/LDH ratio were similar between males and females (CS p=0.949; CS/LDH p=0.271; Fig 3 E and G). When OCR of each mitochondrial complex were normalized to CS activity, males retained higher respiration rates than females (C-I/CS p=0.002; C-II/CS p=0.001; C-III/CS p<0.0001; CIV/CS p<0.001; Fig 4 A-G).

**Fig. 3:**
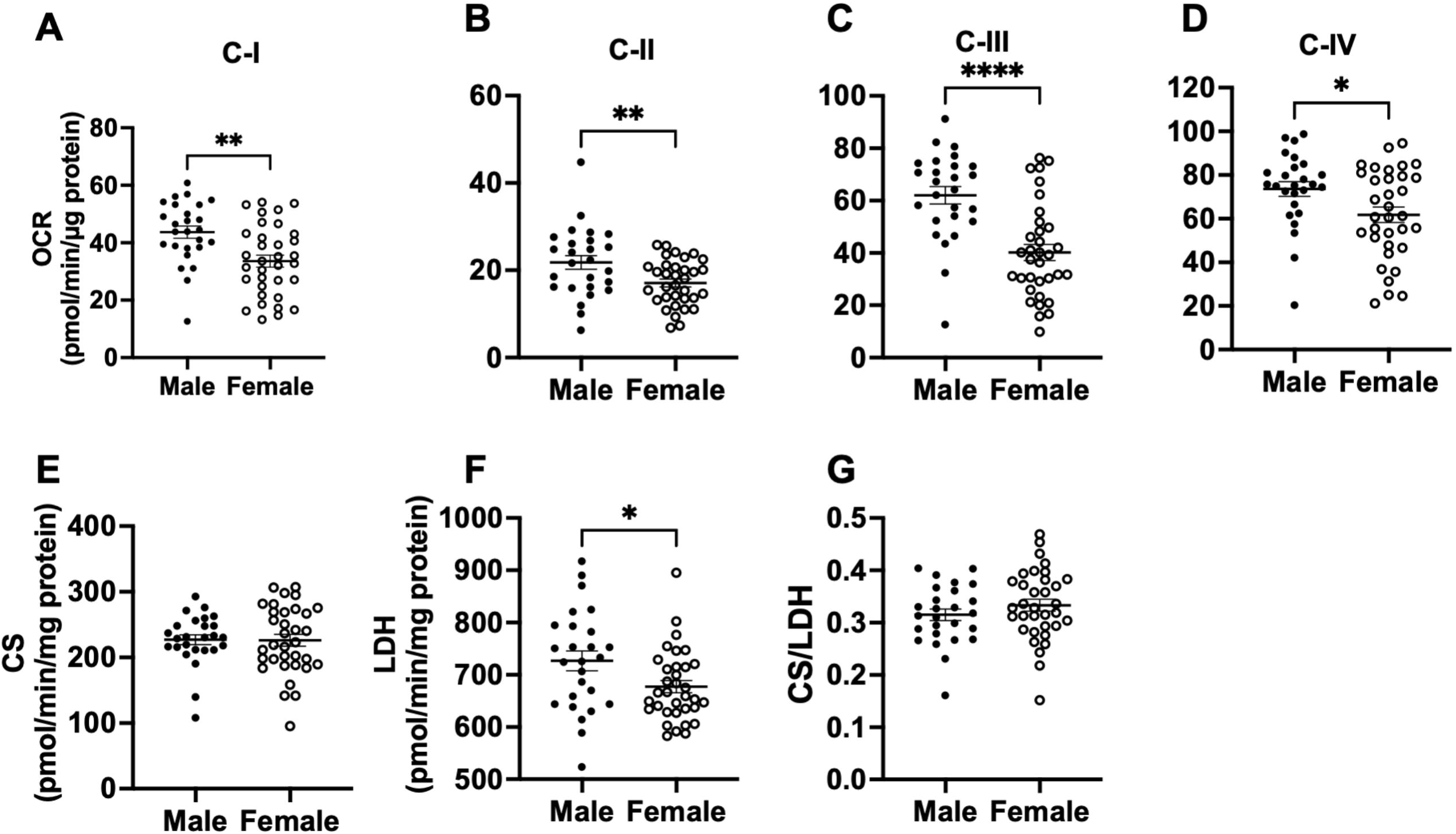
Comparison of mitochondrial complexes I-IV, citrate synthase, and lactate dehydrogenase activities in the prefrontal cortex between male and female baboons. Mitochondrial complex I (C-I) activity (A), C-II activity (B), C-III activity (C), C-IV activity (D), citrate synthase activity – CS (E), lactate dehydrogenase – LDH (F), and LDH to CS ratio (G) between male and females. Each dot represents data from an individual animal, with filled dots representing males (n=24) and open dots representing females (n=36). Error bars indicate mean ± SEM. Statistical significance was determined using t-test, with *p < 0.05, **p < 0.01, and ****p < 0.0001 indicating significance levels.

**Fig. 4:**
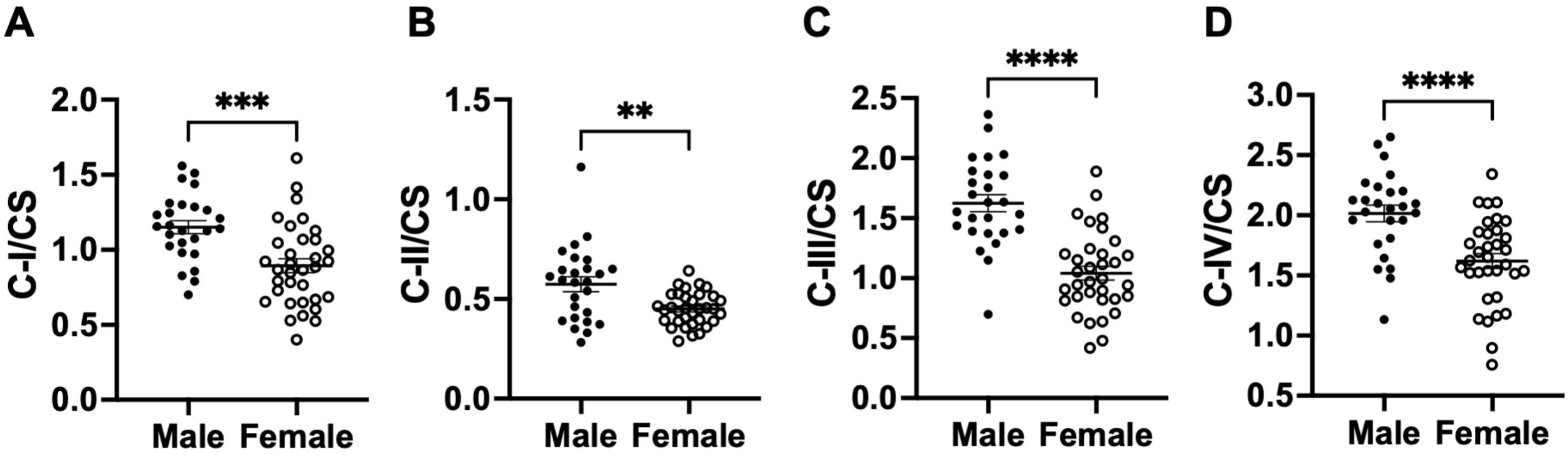
Comparison of normalized mitochondrial activities in the prefrontal cortex between male and female baboons. Comparison between citrate synthase (CS)-normalized mitochondrial complex I: C-I/CS (A), complex II: C-II/CS (B), complex III: C-III/CS (C), and complex IV: C-IV/CS (D) between male and female baboons. Filled dots represents data from males (n=24) and open dots data from females (n=36). Error bars indicate mean ± SEM. Statistical significance was determined using t-test, with **p < 0.01, ***p < 0.001 and ****p < 0.0001 indicating significance levels.

### 3.3. Serum cortisol declines with age in male and female baboons

We confirmed our previous report of an age-related decline in serum cortisol levels in female baboons (Yang et al., 2017) using a larger female cohort in this study (r=-0.51; p=0.018). We observed a similar age-related decline in cortisol levels in males (r=-0.52; p=0.010), and when data from both sexes were combined (Fig 5). Further, there was no correlation between serum cortisol levels and PFC mitochondrial ETC activity in the combined data of male and female baboons (Fig 6 A-D). However, when analyzed by sex, a positive correlation was observed between normalized complex II OCR and cortisol levels in males (p=0.043; Fig 6 F). ETC activity linked to other mitochondrial complexes showed no relationship with cortisol levels in either males or females.

**Fig. 5:**
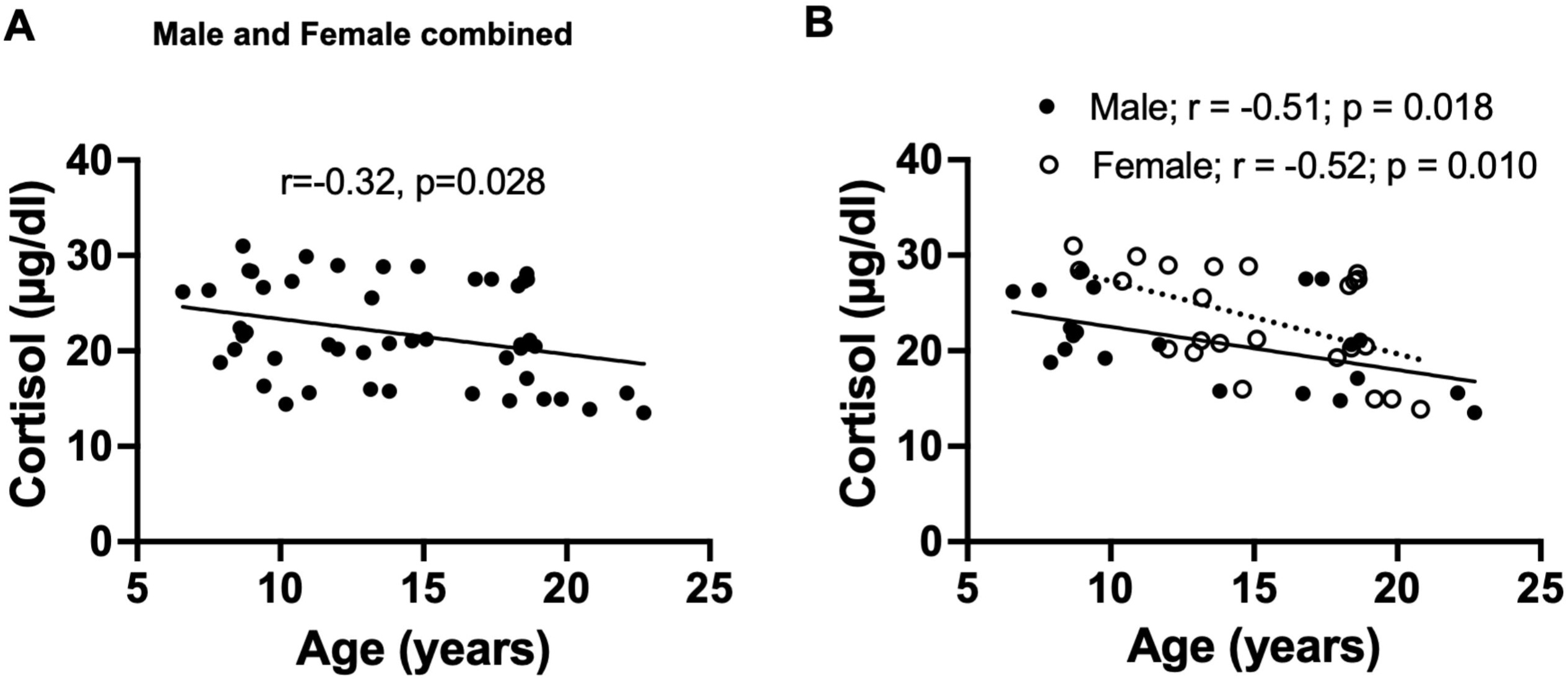
Linear regression of serum cortisol concentration with age in male and female baboons. Plot of serum cortisol concentration versus donor age in combined data of male and female baboons, with regression line represented by a thick line, and each individual dot representing data from an individual animal, n=60 (A). Plot of serum cortisol concentration with age, showing separate analysis for males and females. Filled dots and the thick line depict individual male data and their regression line, respectively, n=24, while open dots and the dashed line represent female data and their regression line, respectively, n=36 (B).

**Fig. 6:**
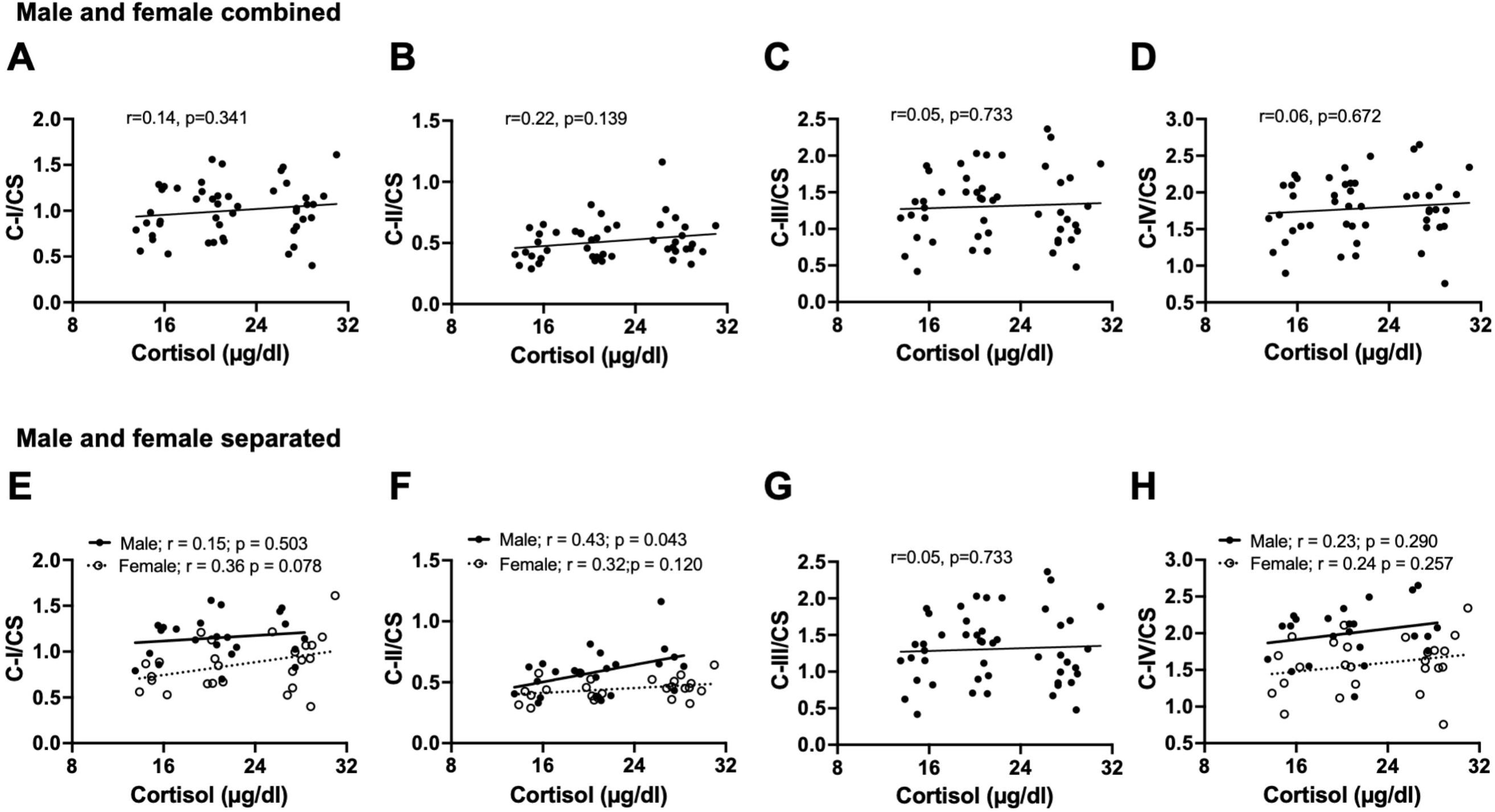
Correlation between circulating cortisol concentrations and normalized mitochondrial activities in the prefrontal cortex of male and female baboons. Pearson’s correlation of serum cortisol concentrations with citrate synthase (CS)-normalized mitochondrial complex I: C-I/CS (A), complex II: C-II/CS (B), complex III: C-III/CS (C), or complex IV: C-IV/CS (D) in combined data from male and female baboons. Regression line represented by a thick line, and each individual filled dot representing an individual animal (n=48). We also separately present sex-specific relationship between serum cortisol and C-I/CS (E), C-II/CS (F), C-III/CS (G), or C-IV/CS (H). Filled dots and the thick line depict individual male data and their regression line, respectively, n=23, while open dots and the dashed line represent female data and their regression line, respectively, n=25.

### 3.4. Male baboon walking speed positively correlates with prefrontal cortex mitochondrial ETC activity

We previously showed that aging is associated with reduced walking speed in both male and female baboons (Huber et al., 2021). We correlated the walking speed data obtained 2 years prior to necropsy with prefrontal cortex mitochondrial ETC activity in the same animal. In the combined dataset of male and female baboons, walking speed positively correlated with citrate synthase-normalized complexes I, II, III, and IV OCR (C-I/CS p=0.001; C-II/CS p=0.032; C-III/CS p<0.0001; C-IV/CS p<0.0001; Fig 7 A-D). We then looked for evidence that there may be sex-specific differences in this relationship with the understanding that these would be limited in power due to the reduced sample sizes. In Fig 7 E-G, we see a positive correlation between CI, CIII, and CIV in males (C-I/CS p=0.033; CII/CS p=0.210; C-III/CS p=0.001; C-IV/CS p=0.001), but not in females (C-I/CS p=0.115; CII/CS p=0.576; C-III/CS p=0.154; C-IV/CS p=0. 0.373), a phenomenon that might be due to a small size or related to slower walking speed in female baboons compared to males.

**Fig. 7:**
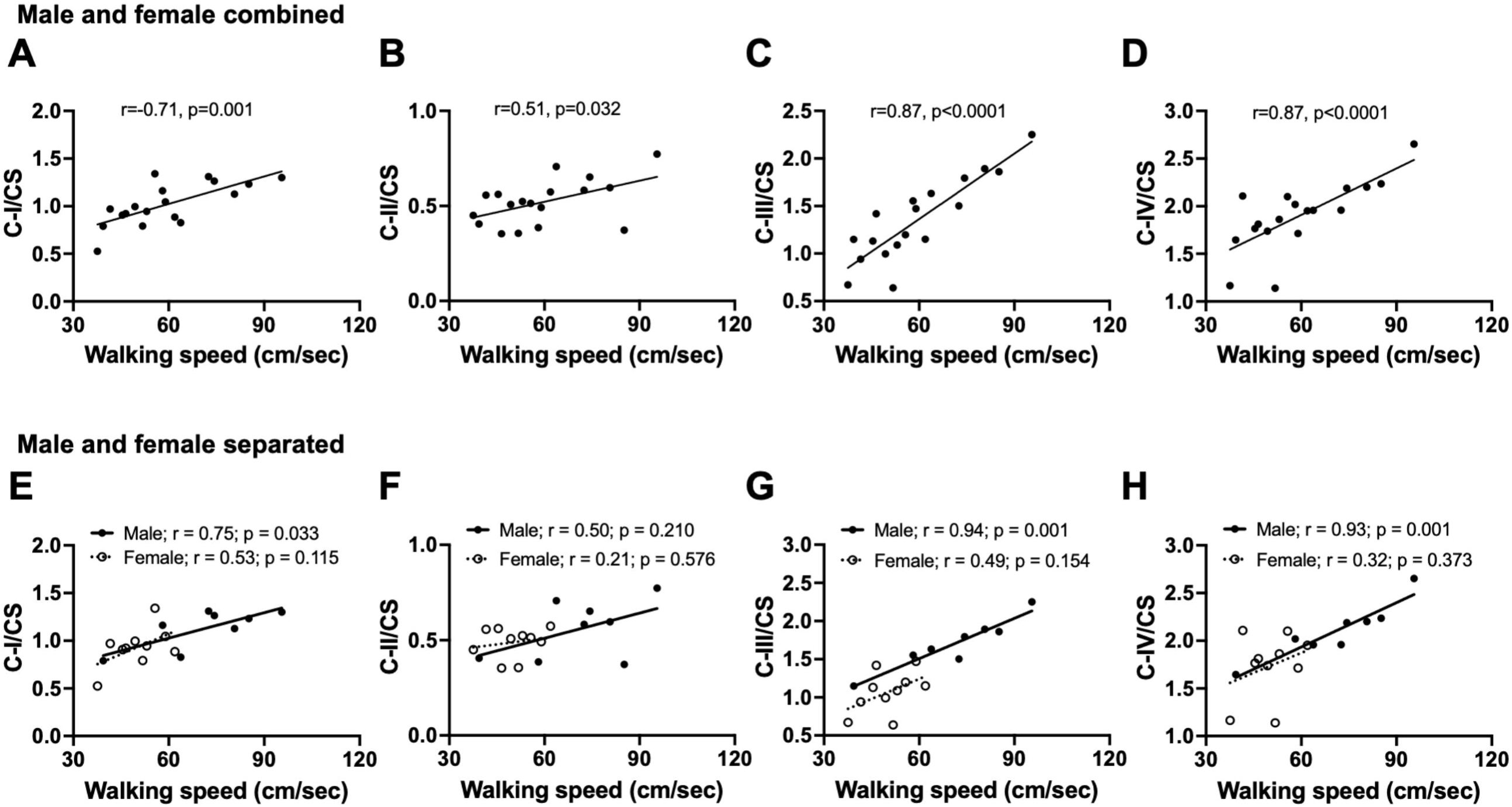
Correlation between walking and normalized mitochondrial activities in the prefrontal cortex of male and female baboons. Pearson’s correlation of walking speed with citrate synthase (CS)-normalized mitochondrial complex I: C-I/CS (A), complex II: C-II/CS (B), complex III: C-III/CS (C), or complex IV: C-IV/CS (D) in combined data from male and female baboons (n=18). Regression line represented by a thick line, and each individual filled dot representing data from an individual animal. Sex-specific relationship between walking speed and C-I/CS (E), C-II/CS (F), C-III/CS (G), or C-IV/CS (H). Filled dots and the thick line depict individual male data and their regression line, respectively, n=8, while open dots and the dashed line represent female data and their regression line, respectively, n=10.

## 4.0. Discussion

This study contributes to the expanding body of evidence that defects in mitochondrial function are a key hallmark of brain aging in a functionally relevant animal model of human aging. Our findings further emphasize that sex is a significant factor regulating differences in the age-related changes in ETC complex activity within the PFC of primates, which may have broader implications for sex-specific resilience mechanisms in the aging brain. Another key finding from this study is that the age-related decrease in respiration linked to ETC complexes positively correlates with walking speed from two years prior, suggesting that walking speed is predictive of decreased PFC mitochondrial bioenergetics, particularly in males. This might also represent a correlation between functional outcome of aging and impact of molecular changes with aging that warrants further exploration. This study also confirms our previous report of an age-related decline in circulating cortisol levels in females, now observed in a larger sample size, and we note a similar age-associated decline in cortisol levels in males, which interestingly correlates with complex II-linked respiration in the PFC.

The human brain is a metabolically expensive organ that uses 20% of the body’s total energy budget despite representing only 2% of total body mass (Raichle et al., 2002). Brain energetics is proportional to neuronal number, with the brain’s energy expenditure directed towards ion pumping to maintain axonal membrane potential and synthesis of neurotransmitters (Herculano-Houzel, 2011, Aiello and Wheeler, 1995). Of the key mitochondrial features that drive mitochondrial function, such as mitochondrial genome, morphology, content, and molecular composition, the ETC is unique for its crucial role in contributing to mitochondrial membrane potential generation and ATP production to fuel cellular processes.

We applied a sensitive high throughput frozen tissue respirometry assay to our archived PFC tissues obtained from a large cohort of baboons, spanning the adult lifespan (∼7-23 years, 60 individuals), to measure maximal respiration linked to ETC complexes I to IV. We observed a clear correlation between donor age and reduced respiration linked to complexes I, II, III, and IV in the PFC. The age-related loss of PFC mitochondrial ETC activity may have implications for cognitive decline with advancing age (Harada et al., 2013). When we tested whether these associations differed based on sex, we found that the age-related decline in mitochondrial respiration was largely driven by males, who exhibited stronger associations than females. Sex-based categorical analysis of mitochondrial respiration independent of age showed that males had higher respiration in the PFC compared to females, a finding similar to higher cortical respiration in adult male mice relative to females (Arias-Reyes et al., 2019). A similar sex-dependent analysis of lactate dehydrogenase (LDH) activity within the PFC showed higher activity in males compared to females, which may imply increased glycolysis in males, given the role of LDH in catalyzing the conversion of glycolysis-derived pyruvate to lactate.

A similar NHP study previously examined age-related changes in mitochondrial respiration in different brain regions of female rhesus monkeys using the Clarke-type oxygen electrode and enzymatic methods on mitochondria fractions (Pandya et al., 2015). The study showed that state III (ADP-stimulated) respiration was significantly reduced with a in hippocampus, substantia nigra, and putamen, and a trend for reduction in the frontal cortex suggesting specific brain region susceptibility to age-related changes in mitochondrial respiration. Mitochondrial enzyme activities of complexes I and IV were unaffected by age in any of the brain regions in this previous study. Our observations in female baboons and the reports above from female rhesus monkeys suggest that the threshold for aging effects on cortical mitochondrial bioenergetics may be higher in females relative to males, which may be due to the influence of female-specific factors such as estrogen in the brain, given that estradiol can regulate mitochondrial bioenergetics (Rettberg et al., 2014). Human studies do not report sex effects, and the combination of male and female data showed a tendency for age-related decline in mitochondrial complex I activity in both the hippocampus and frontal cortex (Yao et al., 2011).

Rodent studies on changes in mitochondrial respiratory rates and enzymatic activities with age show mixed results in males and females compared to primates. One study in only females showed lower activities of complexes I, II, III, and IV in multiple brain regions including the cortex, hippocampus, cerebellum, and brainstem in 24-versus 3-month-old rats (Braidy et al., 2014). A study using only male rats showed that aging decreased mitochondrial enzyme activities, including NADH-cytochrome c reductase (complex I-II) and cytochrome c oxidase (complex IV) in whole brain mitochondria preparations (Navarro and Boveris, 2004). Another male rat study also showed age-related decreases in respiratory fluxes consistent with a decline in complex I activity in brain mitochondria (Cocco et al., 2004). In male mice, enzyme activities of complexes II and III decreased with aging, while complexes I and IV were unchanged (Kwong and Sohal, 1999). A recent mouse study that used a similar respirometry approach to the one we used in baboons showed that respiration dependent on complexes I, II, and IV were lower in the cortex of old male mice relative to young, while in females, only complex II and IV-linked respiration decreased with aging (Sarver et al., 2024). There seems to be a general trend for reduced activity of at least one of the ETC complexes with age in cortical mitochondria of both male and female rodents, whereas primates such as baboons show more robust effects only in males. Species differences, assay variations, and brain region specificity may account for some of these disparities. It is also worth noting that most rodent studies focus on limited age ranges (e.g., 2 months vs. 24 months), whereas our baboon study included data from multiple life stages and used linear regression analysis.

Stress is thought to increase cerebral energy metabolism (Bryan, 1990), a process that reflects increased mitochondrial activity in the brain and suggests interaction between stress factors and mitochondrial metabolism. Glucocorticoid receptors (GR) are expressed in many brain regions, including the frontal cortex (Cook and Wellman, 2004; de Kloet et al., 2005). Glucocorticoids affect mitochondrial oxidative phosphorylation capacity (Picard et al., 2018), and reciprocally, defects in mitochondrial ETC activity are associated with hyperactivation of the hypothalamo-pituitary-adrenal axis (Picard et al., 2015). In men, cortisol exhibits a positive association with brain phosphocreatine to ATP ratios (PCR/ATP), a marker of energy production, but not in women (Mosconi et al., 2024). In the present study, we observed an association between circulating cortisol levels and respiration linked to complex II within the PFC of male baboons but no relationship between cortisol and other complex activities. A cautious interpretation might be that the relationship between PFC complex II-linked respiration and cortisol in the male baboons implies a sex-specific brain metabolic response to stress, such that high cortisol levels in males elicit increased respiration to replenish increased ATP demand associated with stress. This selectivity for complex II is not clear but may be related to several factors, including (i) complex II being encoded by nuclear DNA relative to other ETC complexes encoded by mitochondrial DNA (Rustin and Rötig, 2002), (ii) complex II involvement in regulation of steroid metabolism (Bose et al., 2020), (iii) the uniqueness of complex II as part of both the tricarboxylic acid cycle and the ETC (Rustin and Rötig, 2002; Gnaiger, 2024), and (iv) other biological functions of complex II that transcend respiration, such as control of cell fate and signal transduction (Murphy and Chouchani, 2022; Iverson et al., 2023). The implication of a fall in cortisol level with age observed in this study, similar to earlier reports by us and others (Yang et al., 2017, Willis et al., 2014), in parallel with age-related decline in brain mitochondrial respiration on overall organismal aging is not clear. Whether such a fall in cortisol level is protective or detrimental to the aging process should be further investigated, and it is difficult to speculate since we did not find a strong connection between circulating cortisol and brain bioenergetics. Further studies are required in this area.

Earlier studies have shown the involvement of the PFC in locomotor activity (Hamacher et al., 2015, Nóbrega-Sousa et al., 2020). We showed in this combined dataset from male and female baboons a positive correlation between walking speed and respiration two years later, indicating that slow walking speed is predictive of declining respiration. Further analysis showed that this relationship was largely driven by males, with increased walking speed associated with increased respiration linked to complexes I, III, and IV, but not complex II. This finding is similar to the positive correlation found between basal ganglia mitochondrial state III respiration and locomotor activity of female rhesus monkeys (49). Although we did not measure ATP production in the PFC, which is a limitation of this study, our results and those of others (Pandya et al., 2015) suggest that brain ATP synthesis contributes to the age-related decline in walking speed.

In conclusion, we propose that deficits in energy homeostasis due to decreased mitochondrial ETC activity in the PFC may contribute to cognitive decline with advancing age (Harada et al., 2013). Our findings reveal a sex-dependent decline in PFC mitochondrial respiration with age, predominantly affecting males, which is predicted by decreased walking speed measured ∼ 2 years prior to tissue collections, equivalent to approximately 8 years in humans. These changes in brain bioenergetics with age may provide insights into potential sex-specific resilience mechanisms in the aging brain and identify targets for mitigating age-related motor decline.

## Acknowledgements

This study is jointly coordinated by the San Antonio Nathan Shock Center (NIH P30AG013319) and the UAB Nathan Shock Center (NIH P30AG05886). Baboons in this study were maintained under NIH U19AG057758 (PWN, LAC). The authors acknowledge the administrative and technical support of Karen Moore, Dongbin Xie, and Wenbo Qi. We also acknowledge support from the Southwest National Primate Research Center (SNPRC), which is funded by NIH P51 OD011133, and the San Antonio Claude Pepper Older Americans Independence Center (NIH P30AG044271). Thanks to the staff of SNPRC who support the care of the baboons. Schematic image was created with BioRender.com

