## Supplemental Fig 1 for "Sex-specific decline in prefrontal cortex mitochondrial bioenergetics in aging baboons correlates with walking speed"

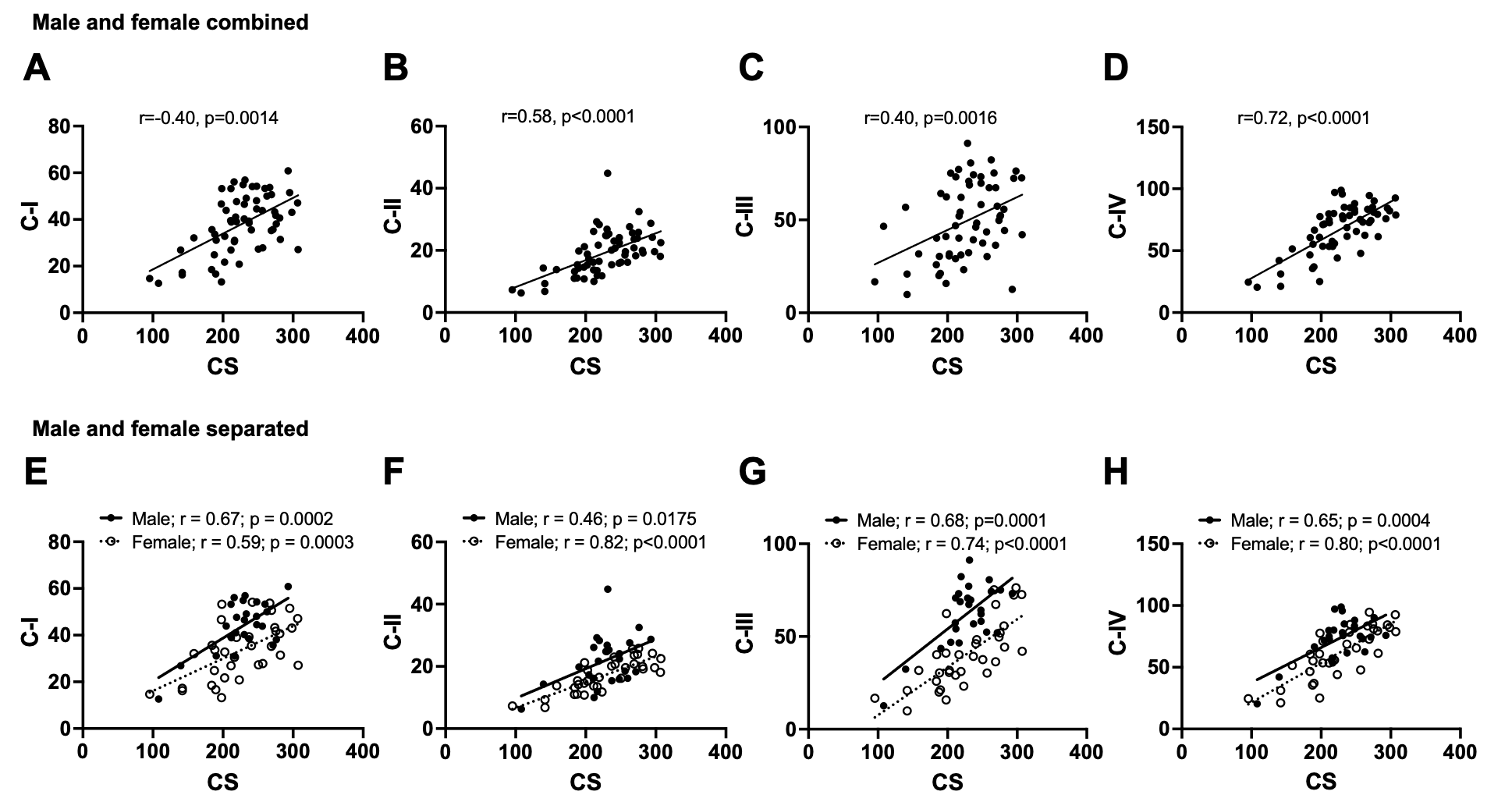

Suppl Fig. 1: Correlation between citrate synthase activity and mitochondrial complexes I-IV activities in the prefrontal cortex of male and female baboons. Pearson’s correlation of citrate synthase activity with mitochondrial complex I: C-I (A), C-II (B), C-III (C), or C-IV (D) in combined data from male and female baboons (n=60). Regression line represented by a thick line, and each filled dot representing data from an individual animal. Sex-specific relationships are shown between citrate synthase activity and C-I (E), C-II (F), C-III/CS (G), or C-IV (H). Filled dots and the thick line depict individual male data and their regression line, respectively, n=24, while open dots and the dashed line represent female data and their regression line, respectively, n=36.
